# Biofilm formation and dynamics in the marine cyanobacterium *Prochlorococcus*

**DOI:** 10.1101/2025.08.05.668435

**Authors:** Maya I. Anjur-Dietrich, Katelyn G. Jones, James I. Mullet, Nhi N. Vo, Kurt G. Castro, Sierra M. Parker, Sallie W. Chisholm

## Abstract

The picocyanobacterium *Prochlorococcus* is responsible for ∼10% of annual marine carbon fixation and plays a role in the global carbon budget. While these phototrophs are primarily considered free-living and neutrally buoyant in the euphotic zone, we observe that they can form biofilms on diverse substrates. This trait is conserved across *Prochlorococcus* ecotypes, and populations continuously transition between planktonic and biofilm states via a non-genetic heritable mechanism. Throughout their growth, cells in biofilms retain a reversible, dynamic attachment state, and measurements of growth, photosynthesis, and respiration rates reveal that cells in biofilms exude more organic carbon than their planktonic counterparts. Estimates of the fraction of *Prochlorococcus* cells attached to particles in the ocean reveal that a significant adherent population exists throughout the euphotic and mesopelagic zones. This work describes a new dimension of *Prochlorococcus*’s ecological niche and suggests a role in carbon export to the deep sea.

## Introduction

Marine phytoplankton account for 49 Gigatons of carbon fixation annually of which nearly 10% can be attributed to a single genus, *Prochlorococcus* ^1^*. Prochlorococcus* is primarily thought of as a neutrally buoyant picocyanobacterium living in the sun-lit euphotic zone, and its importance to marine primary production is firmly established ^1–3^. Because of its efficient turnover in the surface waters, it is thought to contribute relatively little to the ocean’s biological pump, the flux of carbon to, and sequestration in, the deep ocean ^4^. However, *Prochlorococcus* genetic signatures have been found on particles throughout ^5^ and below ^6,7^ the euphotic zone and in mixed-species biofilms grown from seawater ^8^. Further, some *Prochlorococcus* strains are known to attach to and degrade chitin particles ^9^. What is not clear is whether *Prochlorococcus* cells actively form biofilms on surfaces or are simply trapped in the extracellular matrices of other bacteria on sinking particles ^10–13^.

Although much of our understanding of biofilms stems from model organisms or samples collected in medical contexts ^14–22^, it has been argued that most microbes on Earth, whether terrestrial or marine, exist in biofilms ^23^. Basic cellular processes differ between biofilm and planktonic forms, including higher tolerance to antibiotics and environmental stresses such as nutrient starvation and shear ^22,24–27^. In addition, cells in biofilms often have slower growth rates relative to planktonic cells ^28–30^ and may even be dormant or in stationary phase ^31,32^. These differences have revealed new roles for biofilms in nutrient and element cycling in streams ^33^ and soil ^34^, new developmental phases in biofilm formation ^14,19^, and new mechanisms of cell dispersal ^35^.

In *Synechococcus elongatus*, a freshwater genus of cyanobacteria distantly related to marine picocyanobacteria, biofilm formation has been studied in an artificial mutant strain. Wildtype cells self-suppress biofilm formation, and the expression of a biofilm phenotype first requires inactivation of a type IV pilus gene *pilB* ^36–40^. In addition, the self-suppression is caused by a secreted factor, meaning that even Δ*pilB* populations grown in spent media from wildtype populations will not form a biofilm. This result suggests that, in this strain, biofilm and planktonic cells cannot co-exist in the same population, which leaves the role of biofilms in this organism in the natural environment uncertain.

In many non-photosynthetic bacteria, however, the coexistence of—and dynamic transition between—biofilm and planktonic states is a crucial aspect of population survival and spread. In *Pseudomonas aeruginosa*, for example, a cell can become attached within 15 minutes of contacting a surface ^41^, and cells move through states of reversible and irreversible attachment, characterized by the activation of complex regulatory systems ^14,42^. Once in the biofilm, cells can experience varying nutrient gradients and intercellular cues ^31,32,43^, and the transition from biofilm to planktonic, also called biofilm dispersion ^35^, has been linked to responses to these conditions. However, spontaneous dispersal of subpopulations has also been observed ^14,44^, and the precise interplay of attachment and dispersal dynamics, as well as differences between planktonic cells and cells in biofilms, remains an active area of study.

Here, we explore the dynamics and properties of biofilm formation in the marine cyanobacterium *Prochlorococcus*. We used axenic strains representing the breadth of *Prochlorococcus* diversity to investigate the capacity for biofilm formation, established biofilm-forming populations through selection, and then measured physiological and genetic differences between planktonic and adherent cells. We tracked the carbon budget of cells in different states using the rates of photosynthesis, respiration, and cell growth. We further investigated how populations transition between planktonic and biofilm states, and whether selection for either state irreversibly dictates the fate of a population. Finally, we used metagenomic field data to estimate the prevalence and distribution of *Prochlorococcus* cells on particles throughout the global oceans.

## Results

### The biofilm formation trait is found across diverse *Prochlorococcus* clades and is non-genetically heritable

*Prochlorococcus* cells can be broadly classified as high- and low-light adapted ^46,47^–varying in genetic content, nutrient requirements, and light tolerance across latitudes and ocean depths ^48–50^. To systematically explore whether *Prochlorococcus* strains have the capacity to form biofilms and if there are variations among high-and low-light adapted “ecotypes” ^47^, we grew batch cultures of diverse axenic strains representing every clade, hereafter referred to “parental” populations. At mid-exponential phase, populations were separated into two fractions: the liquid fraction, composed of planktonic cells, and the adherent fraction, composed of cells stuck to the walls of the culture tube (Supp. Fig. 1A). The fractions were quantified by either cell counts or chlorophyll *a* absorption with no significant differences between the two methods (Supp. Fig. 1B). Consistently, 6±0.6% (n>=10) of the population in cultures is in the adherent state (Supp. Fig. 1C), and there was no significant difference between high-light and low-light adapted clades (p=0.13) (Supp. Fig. 1C). Thus, *Prochlorococcus* populations have the capacity to form biofilms.

We next explored whether the fraction of cells in either state could shift through selection (Fig. 1A). Starting with mid-exponential parental cultures that had been left undisturbed during growth, an aliquot of the liquid fraction was transferred to new media. After removing the rest of the liquid fraction, fresh media was added to the tube and vortexed to remove any cells adhered to the walls, and an aliquot of the adherent fraction was transferred to new media (Supp. Fig. 1A). This process of serial enrichment, undisturbed growth, and transfer of cells in the adherent state resulted in populations of primarily adherent cells—biofilms—in all *Prochlorococcus* clades examined. Over an average of 26±7 generations, populations transitioned from having 6±0.6% to 85±2% of the cells in the biofilm state (Fig. 1B,C, inset phylogeny adapted from ^51^). High-light and low-light adapted strains reached the same extent of biofilm formation in the population (84±3% and 86±4% adherent cells, respectively, p=0.69). There was no significant difference between high-light and low-light adapted strains in the number of generations required to form persistent biofilm populations (15.2±0.9 generations and 34.8±10.4 generations, respectively, p=0.18), although most high-light adapted strains formed biofilms more readily than most low-light adapted strains (Supp. Table 1). When cells from the liquid fraction of parental strains were selected for and serially transferred, the liquid fraction of the population slightly increased from 92±0.7% to 96±0.3%, leaving approximately 3.8% of the population in the adherent state (Supp. Fig. 1D, final measurement after 120±46 generations across ecotypes).

**Figure 1.**
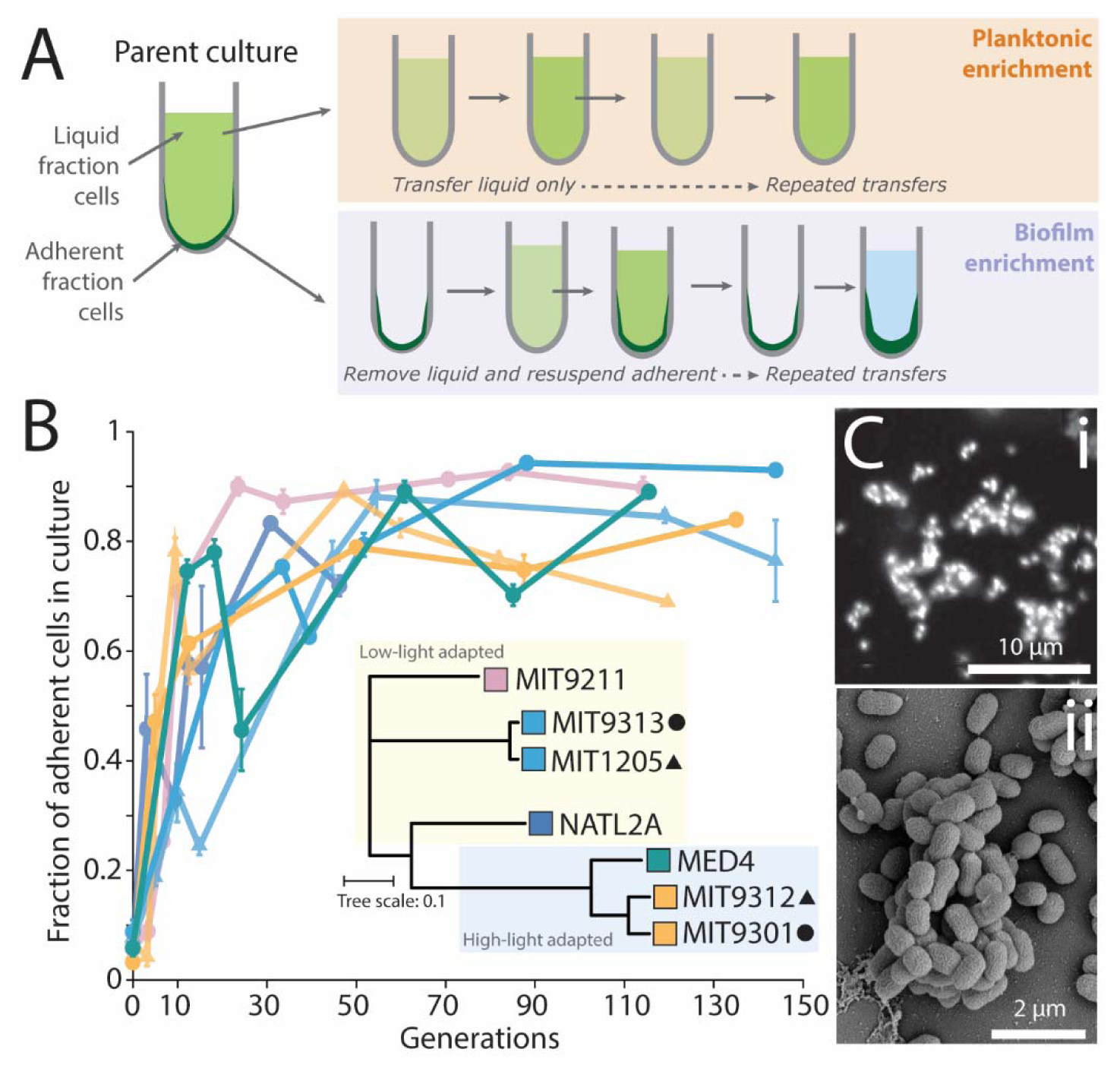
Biofilms can be established in diverse *Prochlorococcus* strains through selection. **(A)** Experimental design of serial culture enrichment of either planktonic or adherent cells. For planktonic enrichments, each culture transfer is started from only planktonic cells. For adherent enrichments, the new culture is started from resuspended (via vortexing) adherent cells. **(B)** Fraction of adherent cells as a function of number of generations of growth in strains drawn from diverse clades of *Prochlorococcus* (inset phylogenetic tree, based on ^51^), all of which start as primarily planktonic parental strains (Supp. Fig. 1). Over an average of 26±7 generations, populations transitioned from 6±0.6% adherent cells to 85±2% with no significant differences between strains from high light-or low-light adapted clades. **(C)** Imaging of cells in biofilms when grown on glass coverslips shown by (i) light microscopy and (ii) electron microscopy.

We also tested whether regularly forcing attached biofilms into suspension without allowing undisturbed growth would cause cells to lose their adherence. We vortexed biofilm cultures daily during growth (as opposed to only when transferring to new media), and, when populations reached late exponential phase, we inoculated new cultures from the vortexed previous culture. Over 30 generations of daily resuspension, cells from biofilm populations re-adhered and passed the adherent property to their daughters (Supp. Fig. 2A,B), indicating that cells from biofilms robustly retain their capacity for adherence.

Thus, adherence and biofilm formation are a phenotype that can be selected for, is transferred across generations, and typically stabilizes when ∼85% of the cells in the population are adherent. Further, the timeline for the transition from a parent to biofilm population is repeatable for a given strain (Supp. Fig. 3A). Cells with the adherent phenotype reattach within 2 days after being disturbed, and the majority of the cells in the new population retain their parents’ adherent properties.

We next investigated whether adherent cells represent a genetically distinct subpopulation in parental strains that were enriched through the selection process. We compared genome sequences of the parental population, the planktonic enrichment, and the biofilm enrichment (each serially selected for over 100 generations) of the high-light MIT 9301 strains using a modified version of the pipeline described in ^52–54^. While we identified SNPs in the parental population relative to the reference genome, these same SNPs were found in all three of the parental, planktonic enrichment, and biofilm enrichment populations (Supp. Fig. 3B). These results are robust as >99% of all reads mapped onto the closed reference genome with an even coverage distribution of reads across the entire genome. Therefore, we conclude that the heritable biofilm formation phenotype is non-genetic, and that individual strains of *Prochlorococcus* have two distinct cell forms—planktonic and adherent—which result in two population states: planktonic and biofilm.

Biofilm formation could be related to a secreted factor, which could be enriched during the serial selection process for adherent cells and underly the increased fraction of adherent cells with each transfer. To test this hypothesis, we grew large volumes of both planktonic and biofilm cultures and either filtered the media with a 0.2 μm filter or centrifuged the media at 6000x*g* to remove cells. We then grew planktonic or adherent cells in both spent media and measured the fractions of cells in the liquid and adherent phases. If adherent cells secreted a molecule that promoted biofilm formation, then planktonic cells would stick when grown in spent media from adherent cells. In addition, if planktonic cells secreted a biofilm-suppressing factor, then adherent cells would fail to form a biofilm when grown in spent media from planktonic cells. In either media condition (filtered or centrifuged), there was no significant difference in the fraction of adherent cells grown in spent planktonic or spent biofilm media or in the fraction of planktonic cells in either media (Supp. Fig. 3C,D). To confirm that the two cell states can stably coexist, we mixed varying percentages of cells from both established planktonic and biofilm enrichments (both selected for over 100 generations), let the cultures grow undisturbed to late exponential phase, and then measured the percentage of cells in the liquid and adherent fractions (Supp. Fig. 3E).

We found that, in both a high-light and low-light adapted strain, the measured culture fractions of planktonic and adherent cells are the same as in the original inoculum after 4 days of growth.

Thus, planktonic cells do not suppress biofilm formation, nor do cells in biofilms cause planktonic cells to adhere or otherwise entrap them.

Together, these results show that biofilm formation is repeatable, non-genetic, and not due to a factor found in spent media. Further, the stable coexistence of planktonic and adherent cells suggests that *Prochlorococcus* biofilms may be a natural state that could be found in wild populations.

### Cells in biofilms have higher photosynthetic rates, lower respiration rates, and slower growth rates than planktonic cells, thus exuding more organic carbon

Growth in a cyanobacterial biofilm can result in self-shading, i.e. the attenuation of light in more basal regions of the biofilm ^55–57^. Indeed, at near-optimal light levels for growth of the parent cells, cells in biofilms grew ∼23% slower than cells from parental cultures in both high-light (MED4) and low-light (MIT9313) adapted strains (Fig. 2A,B). If the growth rate decrease in biofilms were related to self-shading, then an increased incident light intensity would result in an increased growth rate. However, cells in biofilms grew ∼26% slower than parental cells, regardless of light level (Fig. 2A,B, n>=3 and p<0.05 at each timepoint), suggesting light limitation by self-shading is not responsible for the growth rate decrease.

**Figure 2.**
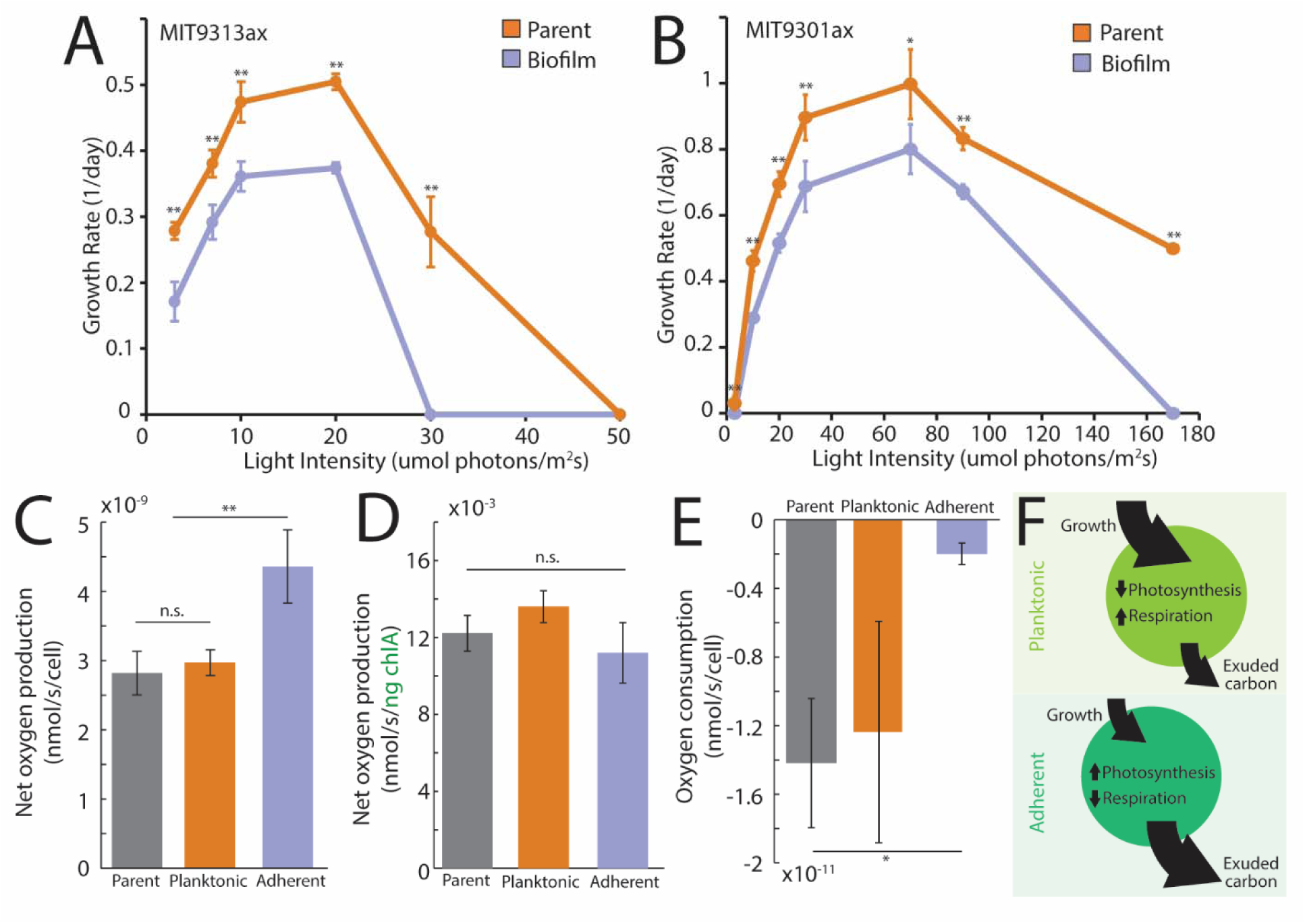
Carbon flux analysis through measurements of photosynthesis, respiration, and growth rates shows that cells in biofilms exude more organic carbon compounds per cell than planktonic cells. **(A)** Growth rate of *Prochlorococcus* cells from parental and biofilm populations of an axenic low-light strain (MIT9313). The biofilm population’s growth rate was 27.9 ± 3.1% slower than the growth rate of the parental populations across all light levels (all biofilm growth rates measured using sacrificial tubes each day). (Significance levels in A and B are * p<0.5, ** p<0.01.) **(B)** Growth rate of *Prochlorococcus* cells from parental and biofilm populations of an axenic high-light strain (MIT9301). Cells in biofilms had a growth rate 25.1 ± 3.0% lower than the parental population across all light intensities. **(C)** Net oxygen production rate per cell in parental, planktonic enrichment, and biofilm enrichment populations (enrichments each selected for over 100 generations). Adherent cells net produced 2.6×10^-10^ nmol O_2_/s/cell, while parental cells net produced 1.6×10^-10^ nmol O_2_/s/cell (n=6, p=0.003). There was no significant difference in the rates of parental and planktonic enrichment cells (planktonic 3.0×10^-9^ nmol O_2_/s/cell, n=6, p=0.69). **(D)** Net oxygen production rate (shown in C) normalized by total chlorophyll *a*. **(E)** Oxygen consumption rate per cell in parental, planktonic enrichment, and biofilm enrichment populations. Parent: 1.42×10^-11^ nmol O_2_/s/cell, n=7; planktonic: 1.24×10^-11^ nmol O_2_/s/cell, n=4; biofilm: 2.0×10^-12^ nmol O_2_/s/cell, n=5. **(F)** Illustration summarizing that planktonic cells have higher growth rates, lower photosynthetic rates, and higher respiration rates than cells in biofilm, i.e. carbon flux balance results in relatively less exuded carbon. Cells in biofilms have lower growth rates, higher photosynthetic rates, and lower respiration rates than planktonic cells, thus exude more carbon than planktonic cells.

Another possibility for decreased growth rates is that cells in biofilms are in a dormant or stationary state ^28,58,59^, as has been previously shown in studies on *E. coli* ^32^ and *P. aeruginosa* ^31^ and as a potential reason for antibiotic resistance in biofilms ^29,60^. To determine whether *Prochlorococcus* adherent cells could be in a depressed metabolic state, we compared the net oxygen production rate of mid-exponential parental and biofilm cultures, which is the gross oxygen production rate minus the respiration rate. Adherent cells net produce 69% more nanomoles of oxygen per second per cell than parental cells (n=6, p=0.03) under the same, near-optimal light and temperature conditions (Fig. 2C). This difference in oxygen production per cell can be explained by increased levels of chlorophyll *a* in adherent cells—adjusting the net oxygen production rate by total chlorophyll *a* removes any significant differences between the three cell populations (Fig. 2D). So, adherent cells have adapted to compensate for self-shading by producing more chlorophyll per cell, which also results in an increased photosynthetic rate. Parental and planktonic cells have indistinguishable net oxygen production rates (Fig. 2A, n=6, p=0.69).

Net oxygen production rate can be larger either because the total rate is higher or because the respiration rate is lower. To distinguish between these two possibilities, we incubated cells in darkness and then measured their cellular respiration rates. We found that cells in biofilms have respiration rates per cell approximately seven times lower than both parental and planktonic cells (Fig. 2E). However, the decreased respiration rate is negligible compared to the oxygen production rate and insufficient to account for the difference in the net oxygen production rate between planktonic and adherent cells, resulting in gross oxygen production by adherent cells approximately 1.5 times higher than parental or planktonic cells (p<0.001). There is no significant difference between gross oxygen production rates of parental and planktonic cells (p=0.25). Therefore, while cells in biofilms have a decreased respiration rate in the dark, they photosynthesize more than planktonic cells under the same conditions.

The balance between photosynthesis and respiration rates determines the carbon available for growth or exudation by a cell. Compared to planktonic cells, adherent cells photosynthesize at higher rates, respire at lower rates, and grow more slowly (all rates in legend of Fig. 2). Therefore, to balance the carbon budget, cells in biofilms must be exuding more organic carbon compounds. This difference suggests that cells in biofilms are likely to create a carbon-rich microenvironment, supporting the growth of surrounding bacteria that feed on organic carbon. It could even provide a way for *Prochlorococcus* cells to build their own particles.

### Adherent and planktonic cells are transcriptionally distinct

We compared the transcriptomes of parental, planktonic enrichment, and biofilm enrichment populations from the high-light adapted strain MIT9301 to explore differences in metabolic state and uncover clues as to the potential selective advantage of cells in a biofilm (Fig. 3, see experimental setup for MIT9301 genome comparison described earlier). Each population was grown to mid-exponential phase in an undisturbed state before collection, i.e. parental and planktonic cells were fixed in suspension, and cells in biofilms were fixed while adhered. Ninety-seven genes were significantly more expressed in cells from biofilms compared to planktonic cells (Fig. 3A, purple shading), and 145 in planktonic cells compared to cells from biofilms (orange shading). Of these genes, two gene sets were represented in their entirety. First, the inorganic phosphate uptake genes (*pstS, A, B, C*) were significantly more expressed in biofilm forming cells, while *phoR*, a negative regulator of the *pst* genes, was more expressed in planktonic cells (Fig. 3A,B). Although this reciprocal pattern seems to suggest that cells in biofilms are phosphorus limited, cultures between 10^5^-10^8^ cells/mL should be phosphorous replete, given the concentrations of phosphate in the media and cell quotas of P ^61,62^. Further, *Prochlorococcus* has been shown to have a low cellular phosphorus requirement ^61–63^ and thrive under very low phosphorus conditions in the oligotrophic oceans ^64^, even substituting S for P in its lipids ^65^. Thus, we explored alternative explanations for the involvement of these genes in biofilm formation. Upregulation of *pstS* in adherent cells relative to planktonic cells has been observed in several *Pseudomonas* strains ^66–68^. As it turns out, *P. aeruginosa* PstS has two distinct domains: one related to phosphate uptake and one related to biofilm formation through an unknown mechanism ^68^. Amino acid alignment between the *Prochlorococcus* MIT9301 and *P. aeruginosa* PstS sequences revealed strong sequence alignment (e-value 8×10^-7^), including in the 15-residue N-terminal region implicated in biofilm formation ^68^ (Supp. Fig. 4). Thus, the upregulation of the *pst* operon in *Prochlorococcus* may play a similar role as in *P. aeruginosa* by contributing to a periplasmic signal that promotes cell adhesion or biofilm formation. However, if the involvement of these genes in the biofilm population also plays a role in P-acquisition, it could be that *Prochlorococcus* biofilms are capable of drawing down phosphorus levels further than planktonic cells, which could affect the relative fitness of adherent and free-living cells in the wild.

**Figure 3.**
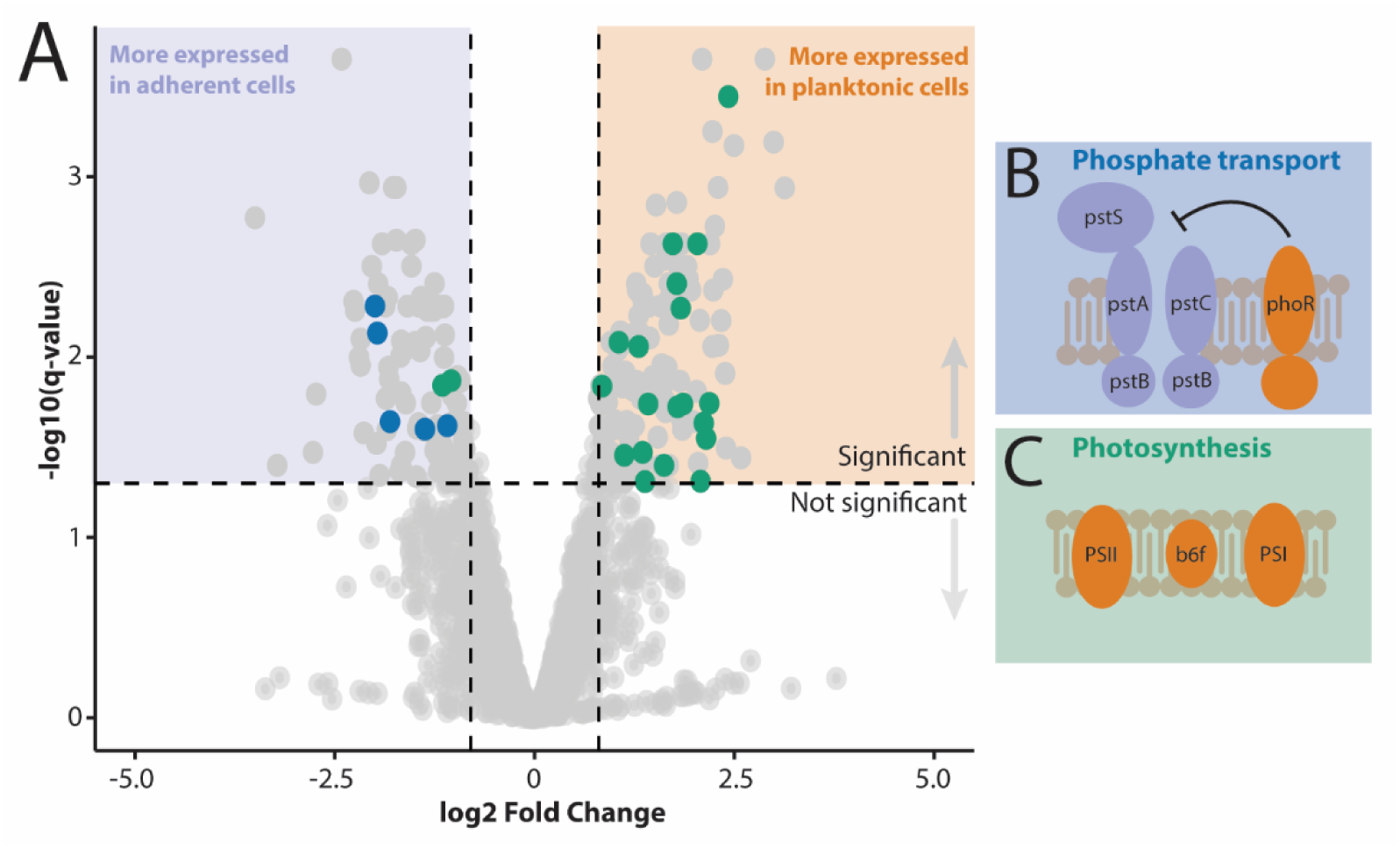
The transcriptional state of cells in biofilms is distinct from that of planktonic cells. **(A)** Fold change (log2) in gene expression between adherent (purple) and planktonic (orange) cells. **(B)** The inorganic phosphate uptake genes *pstS, pstA, pstB,* and *pstC* were more expressed in biofilm-forming cells, while *phoR*, a negative regulator of the *pst* genes, was more expressed in planktonic cells. **(C)** Photosystem and photosynthesis-related genes, including rubisco and chlorophyll synthesis genes, were more expressed in planktonic cells.

Second, all light-dependent photosystem genes and several additional photosynthesis-related genes, including those related to rubisco and chlorophyll synthesis, were expressed significantly more in planktonic cells compared to cells in biofilms (Fig. 3A,C). However, as previously described, this difference in gene expression does not correspond to the relative gross photosynthetic rates. It is possible that relatively higher gene expression reflects a higher protein turnover, which could indicate that planktonic cells more rapidly cycle through photosynthetic machinery, potentially as a result of photodamage. Thus, a decreased photosynthetic protein turnover in adherent cells could also be an effect of self-shading, although further work would be required to confirm this hypothesis.

Homologues of biofilm-related genes that have been identified in several other microbes, including *Vibrio* and *Pseudomonas*, were expressed significantly more in *Prochlorococcus* adherent cells relative to planktonic cells, although no complete pathways were identified. In addition, by using both gene homology searches and protein domain characterization to investigate hypothetical proteins, we identified a homologue of *ebfG*, a small secreted protein required for biofilm formation in *Synechococcus elongatus* ^36^, that was also expressed significantly more in adherent cells. Although the precise interactions of these and other genes related to biofilm formation is still an active area of study, these results suggest that *Prochlorococcus* may use common bacterial biofilm formation pathways.

### Adherent cells emerge from biofilms throughout the course of biofilm growth

Given that cells in biofilms are phenotypically distinct from planktonic cells and transmit these properties to their daughters, we next explored whether being in a biofilm is an endpoint for a cell lineage. We have already shown that repeatedly disturbing (vortexing) adherent cells does not change their capacity to form biofilms, which could suggest that if a cell is adherent, all of its daughters are also adherent and cannot change state. However, the presence of adherent cells in parental populations—and even at low percentages in planktonic-enriched populations—suggests that there must be some rate of spontaneous state change. We envisioned a dynamic population of planktonic and adherent cells that is constantly in transition from one state to another with certain probabilities (Fig. 4A). In this scenario, the planktonic population increases through the growth of planktonic cells or through adherent cells emerging to become planktonic, while it decreases through planktonic cells becoming adherent. Thus, the planktonic population’s measured growth rate is a combination of the cell-specific growth rate of planktonic cells, the rate of emergence of cells from the biofilm (either due to a cell detaching during growth or its daughters detaching during division), and the rate of loss of planktonic cells that spontaneously adhere. Similarly, the biofilm population increases through growth of adhered cells or recruitment of planktonic cells that adhere and decreases through emergence of cells from the biofilm. To distinguish between these net fluxes, we measured the rate of change of either the planktonic or biofilm population comprising a combination of these rates, rather than a cell-specific growth rate. If the planktonic population decreases after the media swap, it indicates that the population’s rate of change is dominated by the adherence rate. However, if the planktonic population increases, the population’s rate of change is primarily due to emergence.

**Figure 4.**
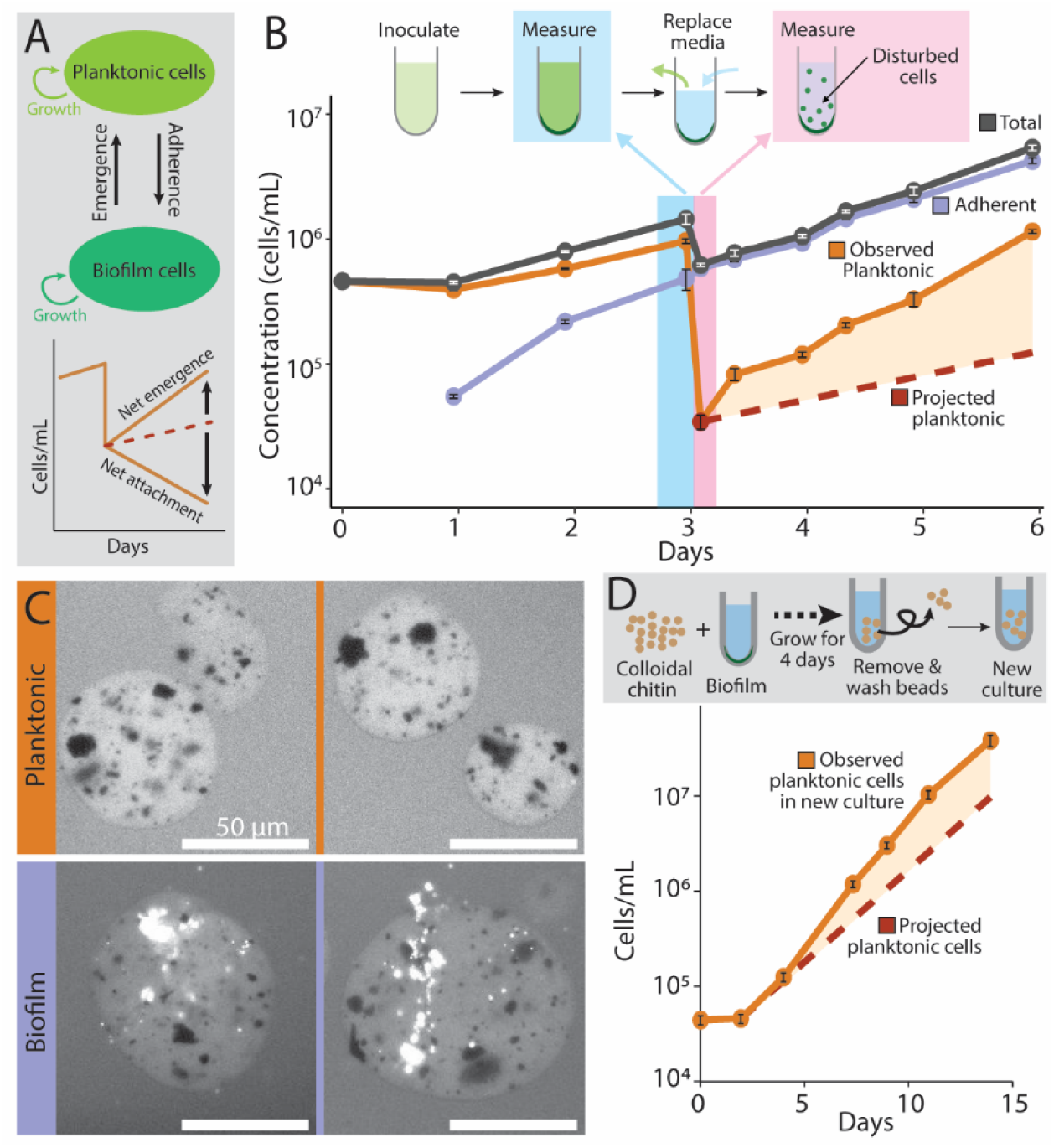
Cells emerge from *Prochlorococcus* biofilms and enter the planktonic phase throughout the growth of the biofilm. **(A)** *Prochlorococcus* populations consist of planktonic and adherent cells with constant transitions between the two states. The planktonic population increases through growth of planktonic cells or through cells emerging from biofilms to resume planktonic growth and decreases through planktonic cells becoming adherent. Similarly, the biofilm population increases through growth or adherence of planktonic cells and decreases through emergence. The rates of change of the planktonic and biofilm populations comprise a combination of these rates, and the relative magnitude determines whether the population undergoes net emergence or net adherence. **(B)** Population growth of adherent cells, planktonic cells, and “newly planktonic”, or cells that emerged from a biofilm. Using the population growth rate from before the media replacement, the projected planktonic population was calculated (red dashed). The planktonic population was ∼65% larger than expected with a significantly increased population growth rate (shaded orange, 0.45±0.021 days^-1^ to 1.15±0.82 days^-1^, p=0.0025, n=3). The growth rate of adherent cells remained similar throughout (days 0-3: 0.86±0.054 days^-1^ and days 3-6: 0.75±0.043 days^-1^, respectively, p=0.17, n=3). The net chance that a cell originating in a biofilm will become planktonic is 38±7% throughout the growth of the population (error is std. dev.). **(C)** Fluorescence images of colloidal chitin magnetic particles (dark spots are magnetite core) incubated with either planktonic (orange) or biofilm (purple) *Prochlorococcus* cells. Cellular fluorescence (bright white) is from chlorophyll (excitation 480/30 nm, emission LP 600 nm). **(D)** Growth of the planktonic fraction of cells in a culture started from chitin particles that were initially incubated with cells from biofilms. The chitin particles were washed several times to remove lightly adhered cells before being transferred to a new culture. The final observed planktonic population exceeded the projected population by 72%, and the net probability of emergence was 44±12% (error is std. dev.).

To determine whether cells in a *Prochlorococcus* biofilm population dynamically adhere and detach and what this means for the overall fate of the population using our framework, we inoculated cultures with cells resuspended from a mid-exponential biofilm and let the population establish itself for 3 days (during which time a robust biofilm population formed), while measuring cell concentrations in the liquid and adherent population fractions each day (Fig. 4B). On day 3, we removed most of the planktonic cells (with a low density remaining to restart the planktonic population) while leaving the biofilm undisturbed (Fig. 4B, pink highlight), added fresh media, and again monitored cell concentrations in the liquid and adherent fractions.

Knowing the planktonic cell concentration when fresh media was added and the net rate of change of the planktonic population in the first 3 days of growth (Fig. 4B, orange line, 0.45±0.021 days^-1^), we could project the trajectory of the free-living population if assuming that increasing cell numbers were largely due to the growth of planktonic cells only (red dashed line days 3-6), since the biofilm population had not yet been established during the first 3 days. Then, differences in the concentration of the planktonic cells from that projection would come from emerging adherent cells as the biofilm matured (orange shading). Indeed, the planktonic cell population was approximately 65% higher than expected based on the population’s projected rate of change before the media swap—reflecting an apparent tripling of the population’s rate of change, which increased to 1.15±0.82 days^-1^ (p=0.0025). This indicates that adherent cells were leaving the biofilm to grow planktonically. Further, the rate of change of the biofilm population was similar before and after the media swap (0.86±0.054 days^-1^ and 0.75±0.043 days^-1^, respectively, p=0.17, n=3), indicating that replacing the media did not affect the division rate of cells in biofilms or the rate at which cells were emerging from the biofilm.

We next characterized the net fate of cells in the biofilm. First, we determined how many more cells are planktonic per day than estimated based on the population growth rate before the media swap, representing the net number of emerging cells over one day. Then we determined the net increase in the biofilm population over the same period, corresponding to a combination of biofilm cell doubling, cell emergence, and cell adherence. Finally, we calculated the percentage of total adherent cells that emerged: (emerged cells)/(emerged cells + biofilm population). This fraction is the net chance that a cell originating in the biofilm will emerge during growth; in other words, the combination of the probability that an adherent daughter cell will emerge from the biofilm minus the probability that a planktonic cell will spontaneously adhere. We found that 38±7% of cells originating in biofilms will become planktonic, and that this percentage is consistent throughout the growth of the population (Fig. 4A). Thus, *Prochlorococcus* populations display an extended period of reversible attachment, and cells continuously emerge from biofilms throughout their growth, not only when it reaches late exponential phase.

In these experiments, cells could emerge from biofilms either by transitioning from an adherent to a planktonic state or by dividing and producing one or two planktonic daughter cells. In *P. aeruginosa*, both reversible and irreversible attachment states have been observed ^14,69^, with the irreversible state often reached in as little as 15 minutes after contacting a surface ^41^. Adherent *Prochlorococcus* cells seem to have a much longer period of reversible attachment, although there may be some heterogeneity in the time an individual cell retains a reversible state. While the net dynamics of adherent and planktonic cells will dictate the fate of overall populations, the attachment and detachment trajectory of individual cells and their daughters would be relevant on shorter timescales.

We next sought to determine patterns of cell emergence on more ecologically relevant substrates, such as chitin, the most abundant marine polymer ^70^ and a substrate that *Prochlorococcus* cells would encounter in the wild. First, we tested whether biofilms would form on colloidal chitin particles. Previous work has suggested that chitin degradation genes are required for the attachment of *Prochlorococcus* cells to chitin particles ^9^, but parental strains were used in that study, which are mainly composed of planktonic cells. We inoculated cultures with cells from planktonic or biofilm MIT9301 enrichments and then added colloidal chitin beads (Fig. 4C). After growing the cultures to mid-exponential phase, there were almost four times as many cells on chitin particles from cultures with adherent cells than from cultures with planktonic cells (9.9×10^4^±2.4×10^3^ cells/mL and 2.5×10^4^±1.6×10^4^ cells/mL, respectively; n=3, Supp. Fig. 5), despite there being 100 times fewer cells in the liquid culture fraction (n=6, p=0.05, Supp. Fig. 5). Thus, it seems that even cells that cannot degrade chitin can attach to this surface as part of a biofilm population.

To test whether adherent cells could detach from chitin beads and then grow planktonically as was observed in glass tubes, we transferred washed chitin beads from cultures with biofilms—i.e. beads with cells adhered—to new cultures and monitored whether any cells appeared in the liquid fraction of the culture (Fig. 4D). Any planktonic cells would have originated from the biofilm on the chitin beads. Planktonic cells in cultures with beads grown with biofilm populations grew at 0.63±0.013 days^-1^, significantly higher than expected based on previous measurements of planktonic MIT9301 cells from biofilm populations (Fig. 4D, n=3, p=0.04). In fact, the final planktonic population in cultures with beads grown with adherent cells exceeded the projection by 72%. Using the framework described above (Fig. 4B), we found that the population dynamics are dominated by emergence from the chitin particle, and that these cells have a 44±12% chance of resuming planktonic growth, similar to the emergence probability seen in experiments with glass test tubes. This emergence rate results in a planktonic population that eventually outnumbers the population growing on the particles by a factor of 30 (1.5×10^6^±5.2×10^5^ cells/mL, n=3), showing that, even with more complex substrates like a chitin bead, biofilm populations maintain an exchange between planktonic and biofilm states. Thus, cells attached to particles in the ocean could either remain attached, ultimately sinking out of the euphotic zone, or eventually detach, setting up the possibility that planktonic populations could be restarted by particle-bound cells.

### Over tens of generations, entire populations dynamically transition between biofilm and planktonic states

Results described above show that selective enrichment produced stable populations: adherent cells reform quickly biofilm populations in new cultures after disturbance, and planktonic cells remain in suspension. Further, neither resuspending cells through vortexing one time nor daily over tens of generations is sufficient to change adherent cells into permanently planktonic (Supp. Fig. 2A, B). However, given that significant numbers of cells spontaneously emerge from biofilms to resume planktonic growth throughout the establishment of a biofilm, it suggests that there are three distinct cell states: planktonic, adherent, and “newly planktonic”, meaning planktonic cells that spontaneously emerged from a biofilm (in contrast to cells that are planktonic after being forced out of the adherent state, which we have already shown will quickly re-adhere and maintain their adherent properties for many generations). The “newly planktonic” cells may have distinct population dynamics and potentially represent a state that more readily transitions between biofilm and planktonic growth, or these cells may behave similarly to planktonic cells in parental cultures.

We tested whether repeatedly selecting only the “newly planktonic” cells from a biofilm population could trigger a shift of the population into a persistent planktonic state or whether these cells have similar dynamics to forcibly disturbed cells which immediately re-adhere (Supp. Fig. 2A,B). At first, inoculating a culture with newly planktonic cells from biofilms resulted in a population with most cells in the adherent fraction (Fig. 5A). Over repeated selections of only “newly planktonic” cells in this population that started from cells in biofilms, the population slowly transitioned away from forming a biofilm and eventually returned to a majority planktonic state (Fig. 5A). This shift back to a mainly planktonic culture took approximately twice as many generations (∼60) as the original transition from planktonic to biofilm (Fig 1A). This result shows that there is a crucial difference between cells emerging from a biofilm state spontaneously and cells forced out of a biofilm state (i.e. vortexed).

**Figure 5.**
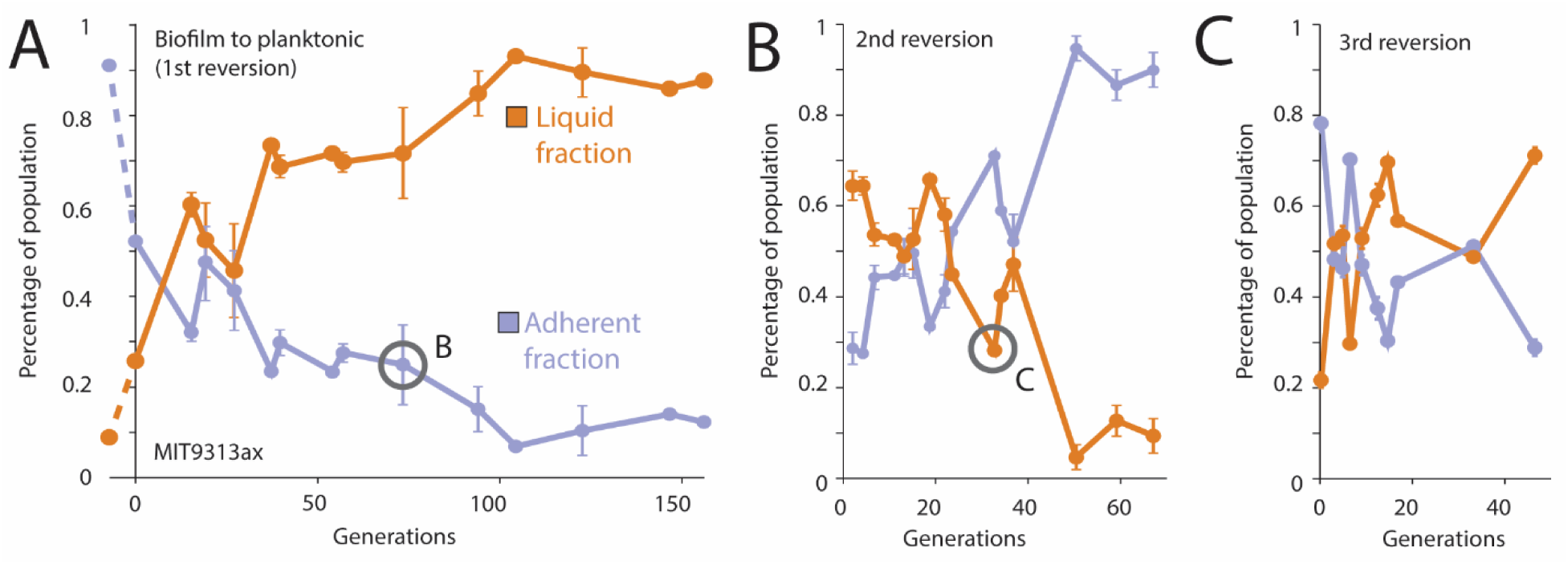
Populations can repeatedly switch between planktonic and biofilm states— showing near-complete reversion of enrichment populations—and these states stably co-exist in culture. **(A)** Fractions of cells in either the liquid or adherent state over time after repeated enrichments of only planktonic cells found in biofilm cultures. **(B)** Cell fractions after repeated enrichments of only adherent cells found in planktonic cultures from time point circled in (A). **(C)** Cell fractions after repeated enrichments of only planktonic cells found in biofilm cultures from timepoint circled in (B).

We repeatedly transitioned populations between planktonic and biofilm states through selection, either taking only planktonic cells from biofilm cultures or adherent cells in planktonic cultures. After showing that newly planktonic cells can eventually form a purely planktonic culture, we next took only adherent cells from this population (grey circle, Fig. 5A) and repeatedly selected for adherent cells (Fig. 5B). We showed that, as with parental populations, we can enrich for adherent cells from a majority planktonic populations. We repeated the transition a third time, selecting newly planktonic cells from the biofilm cultures (grey circle, Fig. 5B) to show that the population could once again become a biofilm (Fig. 5C). We found that repeated transitions do not change the number of generations required to change the state of the population, however, the populations seem to linger in a co-existence state for ∼30 generations before one state rapidly takes over.

### *Prochlorococcus* cells are found on particles from the surface to mesopelagic depths

To better explore the relevance of these results to *Prochlorococcus* in the wild, we examined size-fractionated metagenomic samples collected at Station ALOHA near Hawai’i ^71^ and samples from the global TARA Oceans Project ^72^ (n=579, see Supp. Table. 4 for full sample list). As *Prochlorococcus* cells are less than a micron in diameter, we assume that any cell found in >=1.2 μm fractions must be attached to a particle or other cells. Although the distinction between particle-bound and free-living cells is operational, trends can still be informative. Combining the data from all sites, we see that *Prochlorococcus* cells are found both “free-living” and “particle-bound” (by our definitions) throughout the water column, even well below the euphotic zone (Fig. 6A, Supp. Table 2). While free-living *Prochlorococcus* cells make up a larger fraction of the overall microbial community than particle-bound cells above 150 m, the population fractions are similar at 300 m. At 1000 m, where free-living *Prochlorococcus* makes up less than 1% of the overall microbial community, particle-bound *Prochlorococcus* cells are more prevalent. Below the euphotic zone, the survival of particle-bound *Prochlorococcus* may be supported by the heterotrophic community ^73^.

**Figure 6.**
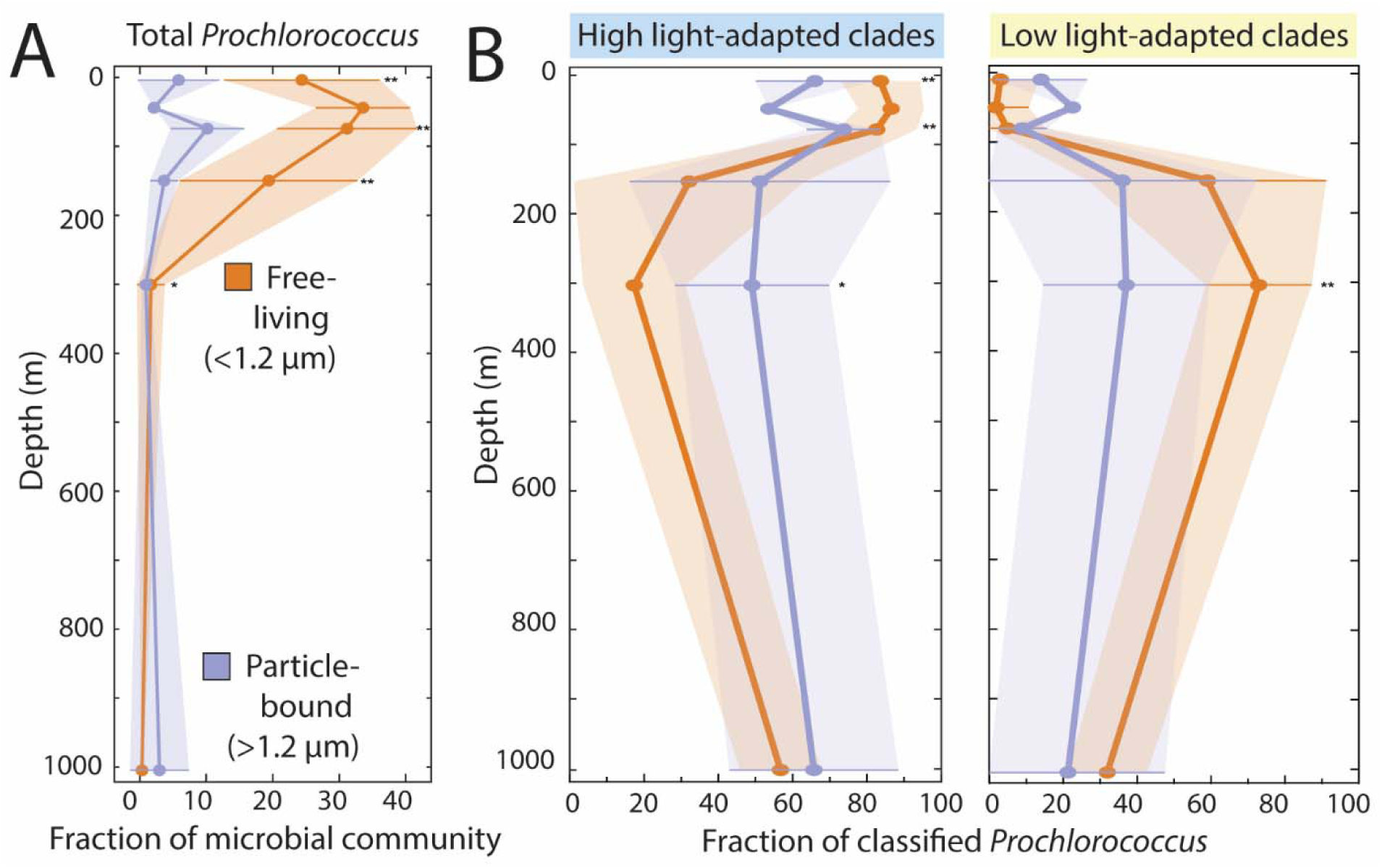
*Prochlorococcus* cells are found both free-living and on particles from the surface to the mesopelagic ocean. **(A)** *Prochlorococcus* genome equivalents as a fraction of the total microbial community in free-living and particle-bound size fractions. (Significance levels in A and B are * p<0.5, ** p<0.01.) **(B)** Clade distribution of *Prochlorococcus* genomes identified in (A).

We examined whether the broadly defined ecotypes (high-light and low-light adapted) were differentially distributed between free living and particle bound fractions as a function of depth. Cells belonging to either clade were generally equally likely to be free-living and particle-bound throughout the euphotic zone, although cells from high-light adapted clades were significantly more likely to be found on particles in the mesopelagic zone (Fig. 6B, Supp. Table 3). The fact that we see ecotype differences between free-living and particle-bound cells as a function of depth lends support to the idea that *Prochlorococcus* cells found on particles are not simply passively trapped from the surrounding water (in which case, population distributions would likely be more similar) but may actively adhere as a response to some condition.

The relative comparisons of free-living and particle-bound population compositions are not sufficient for determining the percentage of the overall *Prochlorococcus* population found either on or off particles, which is the more relevant metric. Thus, we collected a new set of samples from Station ALOHA, to which we could add internal DNA standards, enabling quantitative comparison of fractions of *Prochlorococcus* in either “free-living” or “particle-bound” samples. In the euphotic zone (5-150 m) 11.7±4.7% of *Prochlorococcus* were found on particles (free-living n=27, particle-bound n=25). We calculated the average *Prochlorococcus* cell concentration measured by flow cytometry for HOT cruises 324-339 (^Supp. Table 6^), used this concentration as the 89% planktonic population (since 11% of the total population is on particles), and then calculated what cell concentration would be particle-bound. This fraction translates to approximately 2×10^4^ cells/mL attached to particles at this site. This previously unaccounted for population could play important roles in food web dynamics and carbon export, and future studies should investigate whether the distribution of particle-bound *Prochlorococcus* populations varies across the global oceans.

## Conclusions and Future Directions

*Prochlorococcus* cells are part of a dynamic community in the ocean, comprised of free-living cells and cells attaching and detaching from diverse substrates throughout the water column. Thus, *Prochlorococcus* cells have a complex life trajectory for *Prochlorococcus* cells with a continuous exchange between planktonic and biofilm states (Fig. 7). Planktonic cells—whether those from “wildtype” parental cultures or planktonically-enriched—largely remain planktonic with only ∼6% of cells spontaneously adhering. In contrast, once adhered, cells have multiple possible fates. If cells in biofilms are physically disturbed, ∼80% of cells will re-adhere within 2 days of growth, while ∼20% permanently remain in suspension. However, if biofilms are left to grow undisturbed and reach late exponential phase, the net probability that a cell will leave the biofilm in the first week of growth is ∼40%, suggesting that a spontaneous return to a planktonic state is common in adherent cells.

**Figure 7.**
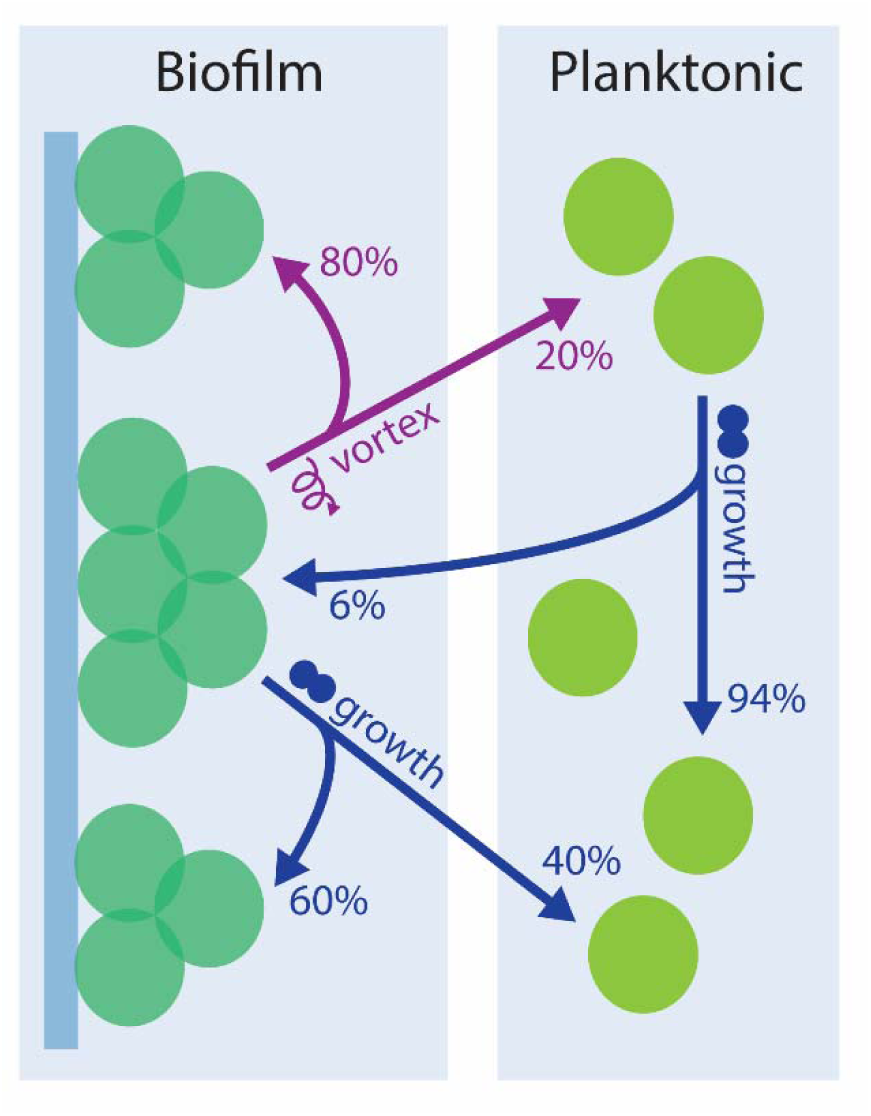
*Prochlorococcus* cells dynamically move between two states, planktonic and adherent, resulting in mixed populations in both biofilm and planktonic states. Approximate transition probabilities between possible cell states are defined, combining population transition data and flow cytometry measurements from Figs. 4 and 5 and Supp. Figs. 1 and 2.

These probabilities describe a population composed of genetically identical cells in multiple phenotypic, heritable states. In other biofilm systems, intercellular variability—or phenotypic diversity—can stem from “microniches”, where cells experience highly localized environments with gradients of nutrients and signaling molecules ^43^ or from stochastic changes in gene expression ^43,74^. In the ocean, particle-bound communities could create their own microenvironment depending on the specific microbes present. The effects of such environments may also be more pronounced in the relatively harsher ocean conditions compared to nutrient-replete lab cultures and could provide a stage for phenotypic diversity to thrive. Even under lab conditions, the observed phenotypic diversity in parental cultures (and selected for in planktonic and biofilm enrichments) clearly results in functional differences between cells, such as differences in growth and photosynthetic rates. Often this type of phenotypic diversity occurs in response to fluctuating environments and represents a bet-hedging strategy for survival, where multiple variants exist in a population when environments change too quickly for individual cells to sense and respond ^74^. In these situations, no interactions between individual cells are required for the evolutionary strategy to pay off. Another possible explanation for phenotypic diversity is division of labor among a population ^74^, which requires some spatial structure in the environment to ensure that all cells have access to any specialized metabolites. While such a strategy would not work for a planktonic *Prochlorococcus* population, the presence of a particle substrate could allow for successful division of labor and a dynamically transitioning population could then extend this survival strategy to planktonic cells.

Attachment and detachment from particles contributes to bacterial population exchange between surface and deeper waters ^75^ and across the global ocean ^76,77^. Although the size and concentration distributions of oceanic particles is uncertain, recent estimates suggest a lower bound of ∼10^5^ particles/mL for particles in the 1-10 μm range ^78,79^. Assuming that individual cells could potentially interact with anything within a radius of 50 μm ^47^, there is a ∼10% chance that any given cell will contact a particle, which would then further increase when taking *Prochlorococcus* concentration into account. Interactions with particles suggest new growth surfaces and dispersal methods for diverse *Prochlorococcus* ecotypes. *Prochlorococcus* cells in biofilms continue photosynthesizing—even shifting their carbon budget to increase exudation— and in sunlit waters could contribute organic carbon to feed particle-associated heterotrophic communities of microbes. As such, it could be a direct contributor of organic carbon to sinking communities and even act as a particle builder, providing initial aggregates to source small particle seeds.

With a global population of ∼10^27^ cells, *Prochlorococcus* is estimated to fix ∼4 Petagrams of carbon annually ^1^. This is on a similar scale to the total amount of carbon exported to the deep sea each year (4-12 Petagrams ^80–84^). Nitrogen tracing of *Prochlorococcus*-derived organic matter has shown that these compounds are incorporated into large, sinking particles ^85^, and, as shown by this work and others, *Prochlorococcus* cells themselves are also found on particles throughout the water column. If 10% of *Prochlorococcus* is attached to particles and fixing carbon as they sink out of the euphotic zone—a reasonable estimate based on our field data—that could move ∼0.5 Petagrams of carbon out of the euphotic zone annually. Given current estimates of carbon sequestration by soft tissue biological pump processes, deep sea carbon export by particle-bound *Prochlorococcus* would be anywhere from 4-12% of the total estimated pump amount, suggesting that *Prochlorococcus* plays a significant and potentially overlooked role ^80–84^. *Prochlorococcus’s* contribution could even increase in the future, as the global population is expected to rise ∼20% by the end of the 21st century ^1,86^, likely including a substantial biofilm population.

## Acknowledgements

The authors would like to thank Easun Arunachalam for valuable discussions. The authors also thank Allison Coe for help with cruise preparation and Alexandrya A. Robinson for experimental assistance. This work was funded by Simons Foundation grants 337262FY23 and 721246. M.A.-D. was supported by the Simons Postdoctoral Fellowship in Marine Microbial Ecology (984601). We also thank the BPF Genomics Core Facility at Harvard Medical School (RRID:SCR_007175) and the BioMicro Center at MIT for sequencing support.

## Author Contributions

M.A.-D. and S.W.C designed the study. M.A.-D. performed experiments, data processing, and statistical analyses with help from K.J., K. C., S.P., and J.M.. N.V. and J.M. performed metagenomic analyses of field samples and mutation analysis with input from M.A.-D. M.A.-D., K.J., S.P., K. C., J.M., and N.V. were supervised by S.W.C. M.A.-D. wrote the paper with input from all authors.

## Methods

### Culturing

All *Prochlorococcus* cultures were grown in 0.2 μm filtered, autoclaved sterile seawater from the Environmental Systems Lab at the Woods Hole Oceanographic Institute amended with Pro99 nutrients as described in ^87^ (Pro99-ESL). Cultures were grown at 24 °C in continuous light near the optimum for each clade (based on data from ^46,49,88^): 35 μmol for MED4 (HLI), 45 μmol photons/m^2^s for 9301 and 9312 (HLII), and 18 μmol photons/m^2^s for NATL2A, 9211, 1205, 9313 (LLI, LLII/III, LLIV, and LLIV, respectively). Cultures were monitored daily by measuring bulk culture fluorescence, and growth rates were calculated by exponential regression from the log-linear portion of the growth curve. Cultures were confirmed as axenic by flow cytometry after staining with SYBR Green (Invitrogen ThermoScientific) for 15 minutes and by inoculating new cultures using broths ProAC, ProMM, and MPTB, as described in ^89–91^.

### Enrichment of planktonic or biofilm populations

All enrichment experiments, meaning any involving selection for a specific culture fraction, were started with an undisturbed, axenic parental culture in mid-exponential phase. For planktonic enrichments, each culture transfer was started from only planktonic cells from the previous culture. For biofilm enrichments, the media and planktonic cells were removed from the parental culture, and the new culture was started from resuspended (via vortexing with fresh Pro99-ESL) adherent cells (method shown in Fig. 1A). At selected timepoints, the percentage of planktonic or biofilm-forming cells in a population was measured as total extracted chlorophyll in methanol from filtered sacrificial samples, which meant the main culture could continue growing undisturbed. Extracted chlorophyll in methanol was measured on a plate reader (Synergy 2, BioTek) and converted to total chlorophyll using absorption coefficients described in ^92^ (method shown in Supp. Fig. 1A). Cell counting by flow cytometry was also used to quantify the percentages of planktonic or adherent cells in a culture. There was no significant difference in culture fractions measured by chlorophyll absorption or cell counting (Supp. Fig. 1B). In experiments referring to the “adherent population fraction”, cells were collected as follows: cultures were grown undisturbed to mid-exponential phase, and then the liquid in the culture tube was removed. Fresh Pro99-ESL was added to the tube and vortexed for 20 seconds to resuspend any cells. In experiments referring to the “liquid population fraction”, cells were collected as follows: cultures were grown undisturbed to mid-exponential phase, and then the liquid in the culture tube was collected.

### Flow cytometry

*Prochlorococcus* cultures were diluted using 0.2 μm filtered Pro99-ESL to at most 300 cells/μL. Cell counts and characteristics (i.e. cell size through forward scatter, chlorophyll content through Red-B) were measured on a Guava flow cytometer (Cytek Biosciences). All flow cytometry runs were started and ended with bead standards to normalize fluorescence intensity across experiments. All flow cytometry data were analyzed using FlowJo version 10.6.1 (FlowJo LLC, BD Life Sciences).

### DNA extraction and sequencing for laboratory experiments

Triplicate biological replicates of cultures were pelleted at 12,000x*g* for 20 minutes. Genomic DNA was isolated using a DNEasy Blood and Tissue Kit (Qiagen). High-throughput libraries were prepared using Nextera FLEX (Illumina) and sequenced with 300 nucleotide paired-end reads using NextSeq 500 (Illumina). SNPs were detected using a modified version workflow described in ^52–54^ with adjusted filtering thresholds (reduced minimum allele frequency threshold of 0.1; maximum reads at a given positions of 5000) (Modified workflow available at https://github.com/nhinvo/WideVariant-SnakemakeOnly). Reads were aligned to the *Prochlorococcus marinus str.* MIT 9301 reference genome (NC_009091.1) with >99% of reads mapping across all samples. Read coverage was between 800X-1000X at most positions with the lowest coverage at ∼400X, and there was even coverage across the entire genome for all samples.

### Field sample collection and metagenomic analysis

Two types of data from samples collected from the oceans were analyzed for this paper: 1) samples collected recently from Station ALOHA in collaboration with the Hawai’i Ocean Time-series (HOT) to which internal standards were added, and 2) reanalyzed samples collected through the TARA Oceans Project ^93^ or at HOT ^71^ (full sample list in Supp. Table 4). Metagenomic samples were defined as either containing free-living microbes, collected from serial filters with pore sizes >0.2 μm and <1.2 μm, or particle-bound microbes, from serial filters with pore sizes >=1.2 μm. Field samples were collected on the HOT346 cruise with the Hawai’i Ocean Timeseries as follows: 500 mL of water collected from 5, 75, 150, 300, and 1000 m was serially filtered through 20, 5, 1.2, and 0.2 μm filters, which were then frozen. DNA was extracted from filters through an adapted phenol-chloroform extraction protocol described in ^94,95^. A fraction of filters was combined and extracted together only if each filter was an identical replicate of the same filter size and depth to maximize DNA recovery. A DNA standard (*Thermus thermophilus* HB27, ATCC BAA-163D-S) was added according to estimated DNA yields at each depth. High-throughput libraries were prepared using Nextera FLEX (Illumina) and sequenced with 300 nucleotide paired-end reads using NextSeq 500 (Illumina). Trimmed and quality-controlled reads and adapters were mapped to the *Thermus* reference genome to identify and remove internal standard sequences. Field samples from the TARA Oceans Project were selected as follows: images with identifiable particles were extracted from EcoTaxa ^96^, and their unique identifiers were matched with metagenomic samples. The filtered reads were taxonomically classified using the ProSynTax workflow ^51^. All classification results were normalized by correcting for differences in average genome length between different *Prochlorococcus* clades. For absolute genome equivalent measurements, the sequencing efficiency per sample was obtained by comparing standard molecules added against standard molecules recovered (the latter estimated by aligning all metagenomic reads to the *Thermus* reference genome using BLAST ^97^. Efficiencies were used to correct the normalized genome equivalent outputs from the ProSynTax workflow. The *Prochlorococcus* cell concentration at Station ALOHA was estimated by calculating the average concentration measured by flow cytometry on HOT cruises 324-339 for 5-150 m (Supp. Table 6).

### RNA extraction and analysis

Triplicate biological replicates of cultures were pelleted at 12,000x*g* for 20 minutes. RNA was isolated using a standard acidic phenol:chloroform protocol, and samples were cleaned using RNAClean XP beads (A63987, Beckman). Libraries were prepared using a KAPA HyperPrep with Ribo-Erase kit (Roche), substituting bacterial NEBNext rRNA depletion probes (New England Biolabs), and then residual primers were cleaned using KAPA Pure Beads in a 0.63x SPRI-based cleanup (Roche). High-throughput libraries were prepared using Nextera FLEX (Illumina) and sequenced with 300 nucleotide paired-end reads using NextSeq 500 (Illumina). Low-quality reads and adapter sequences were moved using BBDuk (v38.16) (minlen=25, qtrim=rl, trimq=10, maq=20, ktrim=r, k=23, mink=11, and hdist=1). Trimmed reads were aligned to the Prochlorococcus MIT9301 genome (NC_009091.1) using bowtie2 (v2.5.4) with default settings. The number of reads that aligned to each annotated open reading frame were counted using HTSeq (v0.11.2) (-s reverse, -t exon, -r pos, –nonunique all). Differential gene expression analysis was performed using edgeR ^98^, setting the log-fold change threshold to 0.8 and the q-value threshold to 0.05. Genes with a log2-fold change less than -0.8 and a q-value less than 0.05 were marked as significantly more expressed in cells in biofilms than planktonic cells. Genes with a log2-fold change greater than 0.8 and a q-value less than 0.05 were marked as significantly more expressed in planktonic cells compared to cells in biofilms.

### Homologous gene identification

The MIT9301 genome was annotated with UniProt. Further searches used the following methods: Identified homologous genes of interest were aligned using T-Coffee ^99,100^ and NCBI Protein Blast ^101^ and visualized with Jalview ^102^. Searches of hypothetical genes began with PSIBlast with 4 iterations across a database of *Prochlorococcus* genomes based on protein sequences from previously described genes in *Synechococcus* PCC7942. The filtering thresholds were set to e-value <1e-5, query coverage (%) > 40, and percent identity > 0.25. For identification of *pteB*, an HMMsearch ^103^ of all proteins in a given strain for ABC transporter and C39 peptidase domains was used. Candidate proteins were inspected using InterPro ^104^, and final candidates were verified by PSIBlast. *ebfG* genes were identified by proximity to *pteB* candidates and verified by PSIBlast.

### Oxygen measurements

Oxygen production rates were measured using an O_2_ UniAmp system (Unisense) equipped with a low range, 500 μm probe. The system was calibrated with a two-point calibration using Pro99-ESL acclimatized to the experiment temperature for at least 90 minutes, measuring the O_2_-replete point, then sparging with CO_2_ for the 0 point. Measurements were performed in a temperature-controlled water bath with calibrated light levels, keeping the same conditions as the near-optimal growth conditions. For axenic 9301 strains, the light level was 32 μmol photons/m^2^s and the water bath was at 24.5 C. Sparged media was used to dilute cultures immediately before measurements, and then oxygen levels were recorded for 10-15 minutes. Rates were calculated from linear regions of the trace, eliminating early adjustment jitter from the probe. Oxygen consumption rates were measured similarly, except that cultures were incubated in darkness for 30-60 minutes before measurements. While in darkness, cultures were placed in the water bath maintained at 24.5 C, and measurements were performed in darkness. Rates were calculated from linearly decreasing regions of the trace, eliminating early adjustment jitter as well as any remnant oxygen production signal.

### Statistical analysis

P-values were calculated using 2-tailed t-tests, assuming equal variances. Unless otherwise specified, all reported errors are standard error of the mean. In cases where sample sizes were significantly different (such as with cruise data), p-values were bootstrapped using the average of 10 subsamples of the original dataset.

