## Supplementary Information for "Biofilm formation and dynamics in the marine cyanobacterium *Prochlorococcus*"

**Supplementary Information (Anjur-Dietrich, *et al.*).**

**Supplementary Table 1. Time in generations to biofilm enrichments in *Prochlorococcus* strains.**  Ecotype abbreviations: HL = High-light adapted, LL = low-light adapted.

| **Strain** | **Ecotype** | **Generations to biofilm** |
| --- | --- | --- |
| MED4 | HL | 17.8 |
| 9301 | HL | 16.1 |
| 9312 | HL | 9.7 |
| 1205 | LL | 51.6 |
| 9313 | LL | 35.0 |
| 9211 | LL | 12.1 |
| NATL2A | LL | 23.0 |
| SB | LL | 13.1 |

**Supplementary Table 2. *Prochlorococcus* fractions of the total microbial community, measured by normalized metagenomic reads.** The aggregated metagenomic collection includes samples from HOT346 (collected for this study), TARA Oceans, and samples described by Leu, *et al.* 2022 from the Hawai’i Ocean Time series (see Supp. Table 4 for identifiers). We define “free-living” as cells collected on filters with pore sizes >0.2 μm and <1.2 μm and “particle-bound” as >=1.2 μm. The combined euphotic zone is defined as 5-150 m, while the combined mesopelagic zone is defined as 300-1000 m. The datasets were subsampled 10 times (choosing the same number of samples from the larger dataset as in the smaller dataset without replacement), and then the p-values were averaged. Errors are reported as standard deviation.

| **Depth (m)** | **Free-living (%)** | **Particle-bound (%)** | **p-value** | **N, Free-living** | **N, Particle-bound** |
| --- | --- | --- | --- | --- | --- |
| 5 | 63±20 | 12±14 | <<0.001 | 67 | 28 |
| 45 | 74±12 | 3±0 | n/a | 71 | 1 |
| 75 | 70±13 | 15±7 | <<0.001 | 42 | 16 |
| 150 | 46±24 | 5±3 | <<0.001 | 108 | 21 |
| 300 | 5±6 | 1±0.9 | 0.04±0.04 | 102 | 11 |
| 1000 | 1±0.3 | 3±5 | 0.3±0.02 (n.s.) | 93 | 10 |
| Euphotic | 60±22 | 11±11 | <<0.001 | 288 | 66 |
| Mesopelagic | 3±5 | 2±3 | 0.4±0.2 (n.s.) | 195 | 21 |

**Supplementary Table 3. Clade breakdown of *Prochlorococcus* populations reported in Supp. Table 1.** HL = High-light adapted, LL = low-light adapted. Average p-values were calculated using 10 subsamples from the larger dataset, choosing the same number of samples as in the smaller dataset without replacement. Errors are reported as standard deviation.

| **Clade** | **Depth (m)** | **Free-living (%)** | **Particle-bound (%)** | **p-value** | **N, Free-living** | **N, Particle-bound** |
| --- | --- | --- | --- | --- | --- | --- |
| HL | 5 | 84±11 | 70±15 | 0.002±0.004 | 67 | 28 |
| HL | 45 | 87±9 | 54±0 | n/a | 71 | 1 |
| HL | 75 | 83±11 | 75±9 | 0.04±0.06 | 42 | 16 |
| HL | 150 | 33±31 | 42±36 | 0.4±0.3 (n.s.) | 108 | 21 |
| HL | 300 | 17±13 | 43±21 | 0.01±0.01 | 96 | 10 |
| HL | 1000 | 68±9 | 62±22 | 0.4±0.09 (n.s.) | 19 | 9 |
| HL | Euphotic | 65±33 | 62±27 | 0.4±0.3 (n.s.) | 288 | 66 |
| HL | Mesopelagic | 25±23 | 52±23 | 0.005±0.008 | 115 | 19 |
| LL | 5 | 3±11 | 10±12 | 0.1±0.1 (n.s.) | 67 | 28 |
| LL | 45 | 2±9 | 23±0 | n/a | 71 | 1 |
| LL | 75 | 5±11 | 8±6 | 0.3±0.3 (n.s.) | 42 | 16 |
| LL | 150 | 59±33 | 46±37 | 0.3±0.2 (n.s.) | 108 | 21 |
| LL | 300 | 74±14 | 44±22 | 0.005±0.0009 | 96 | 10 |
| LL | 1000 | 21±9 | 26±26 | 0.5±0.2 (n.s.) | 19 | 9 |
| LL | Euphotic | 24±34 | 21±28 | 0.6±0.2 (n.s.) | 288 | 19 |
| LL | Mesopelagic | 65±24 | 36±25 | 0.002±0.002 | 115 | 19 |

**Supplementary Table 4. Sample identifiers used in metagenomic analysis of field samples.** Provided as an Excel sheet.

**Supplementary Table 5. Full RNASeq results for transcriptome comparison of MIT9301 planktonic enrichment and biofilm enrichment populations.** Each enrichment was selected for over 100 generations. Provided as an Excel sheet.

**Supplementary Table 6. Raw *Prochlorococcus* cell count data measured by flow cytometry from HOT cruises 324-339.** Provided as an Excel sheet. Data extracted using Hawai'i Ocean Time-series Data Organization and Graphical System.

**Supplementary Figure 1. Parental *Prochlorococcus* populations consistently have a small percentage of adherent cells, whether measured by cell number or chlorophyll. (A)** At each timepoint, the liquid fraction (i.e. media and planktonic cells) was removed and filtered, and then the filter was incubated in methanol. Fresh methanol was added to the original culture tube and vortexed to dissolve any adherent cells. Both samples were incubated at 4 C overnight, and then the percentage of planktonic and adherent cells (in the liquid and adherent fractions, respectively) was measured as total extracted chlorophyll using standard techniques. **(B)** Population fractions of liquid and adherent *Prochlorococcus* measured by chlorophyll absorption or cell counting. The percentage of adherent cells in parental cultures of MIT9313 was 2.0 ± 1.0% measured by chlorophyll absorption and 1.4 ± 0.00015% measured by flow cytometry (n=3, p=0.66). In biofilm populations, 90.4 ± 3.6% of cells were adherent measured by chlorophyll absorption and 95.7 ± 1.0% by flow cytometry (n=3, p=0.31). **(C)** Liquid and adherent population fractions in parental axenic cultures. Across clades, parental strains were composed of 92 ± 0.7% planktonic cells and 6 ± 0.6% biofilm cells (n>= 10 for all strains). There was no significant difference between the fraction of adherent cells in high-light and low-light adapted clades (5 ± 0.9% and 7 ± 0.6%, respectively, p=0.13). **(D)** Population fractions of liquid and adherent *Prochlorococcus* in a planktonic enrichment of a low-light strain (MIT9313).

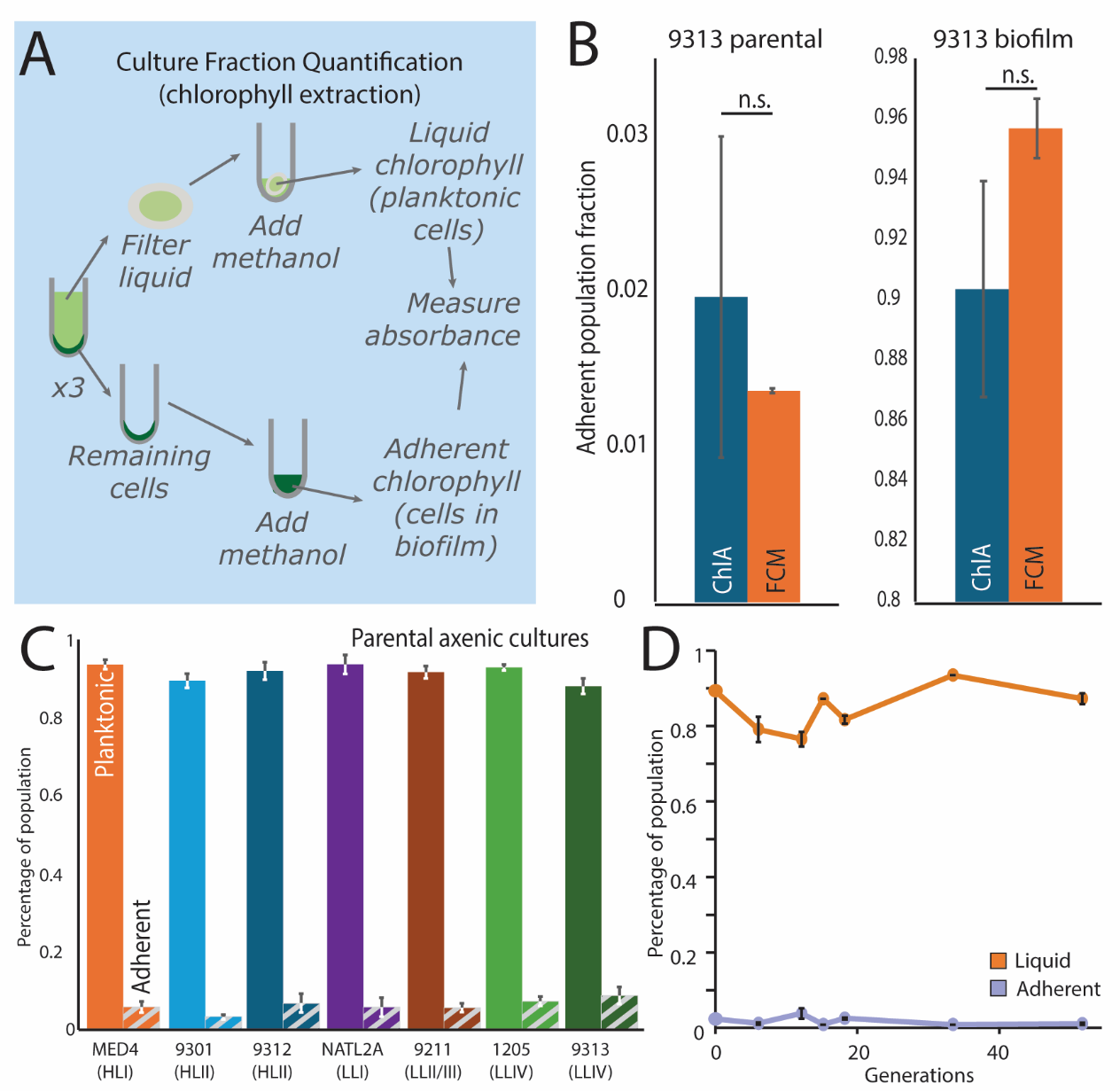

**Supplementary Figure 2. Daily physical perturbation of biofilms is insufficient to return cultures to a planktonic state. (A)** Fractions of a biofilm population from a low-light adapted strain (MIT9313ax) that was vortexed daily. At each transfer, the entire culture was resuspended (rather than isolating only adherent cells as during biofilm enrichment) and used to inoculate a new culture. **(B)** Fractions of a biofilm population from a high-light adapted strain (MED4ax) that was vortexed daily (same experimental setup as described in A).

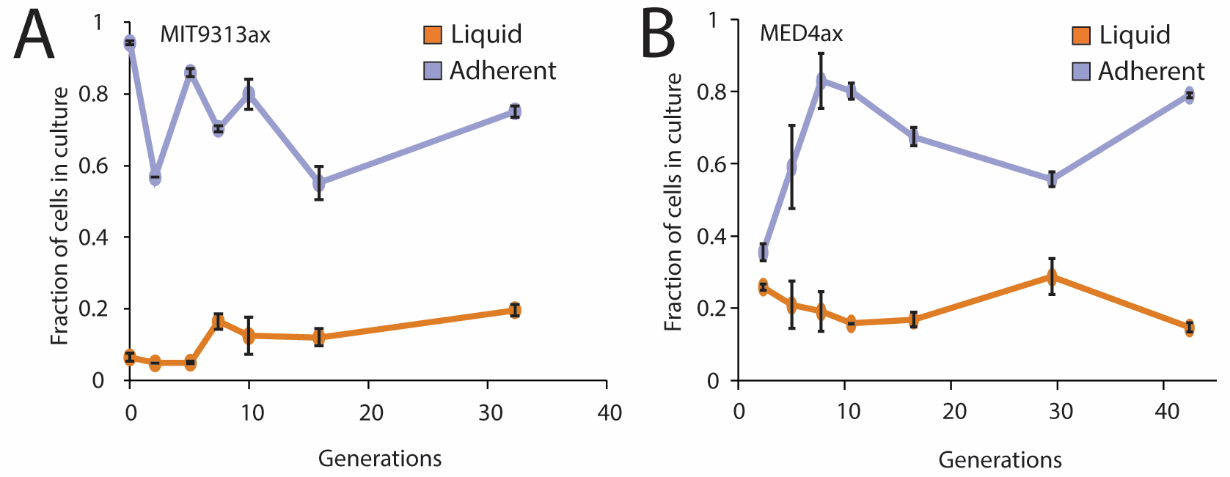

**Supplementary Figure 3. Biofilm formation is repeatable, non-genetic, and not due to a factor found in spent media. (A)** Population fractions measured by chlorophyll *a* absorption from replicate enrichments for biofilms. **(B)** Base calls from a representative position of parental, planktonic enrichment, and biofilm enrichment populations of a high-light strain (MIT9301) mapped onto a closed reference genome (enrichments each selected for over 100 generations). **(C)** Fraction of planktonic and adherent cells in cultures with cells either from planktonic or biofilm cultures grown in centrifuged, cell-free spent media from either planktonic or biofilm cultures (MIT9301). **(D)** Fraction of planktonic and adherent cells in cultures using the same experimental setup as in (C) with the exception that the spent media was 0.2 μm-filtered (MIT9301). **(E)** Initial inoculation ratio of adherent to planktonic cells compared to measured adherent fraction after 3 days of growth.

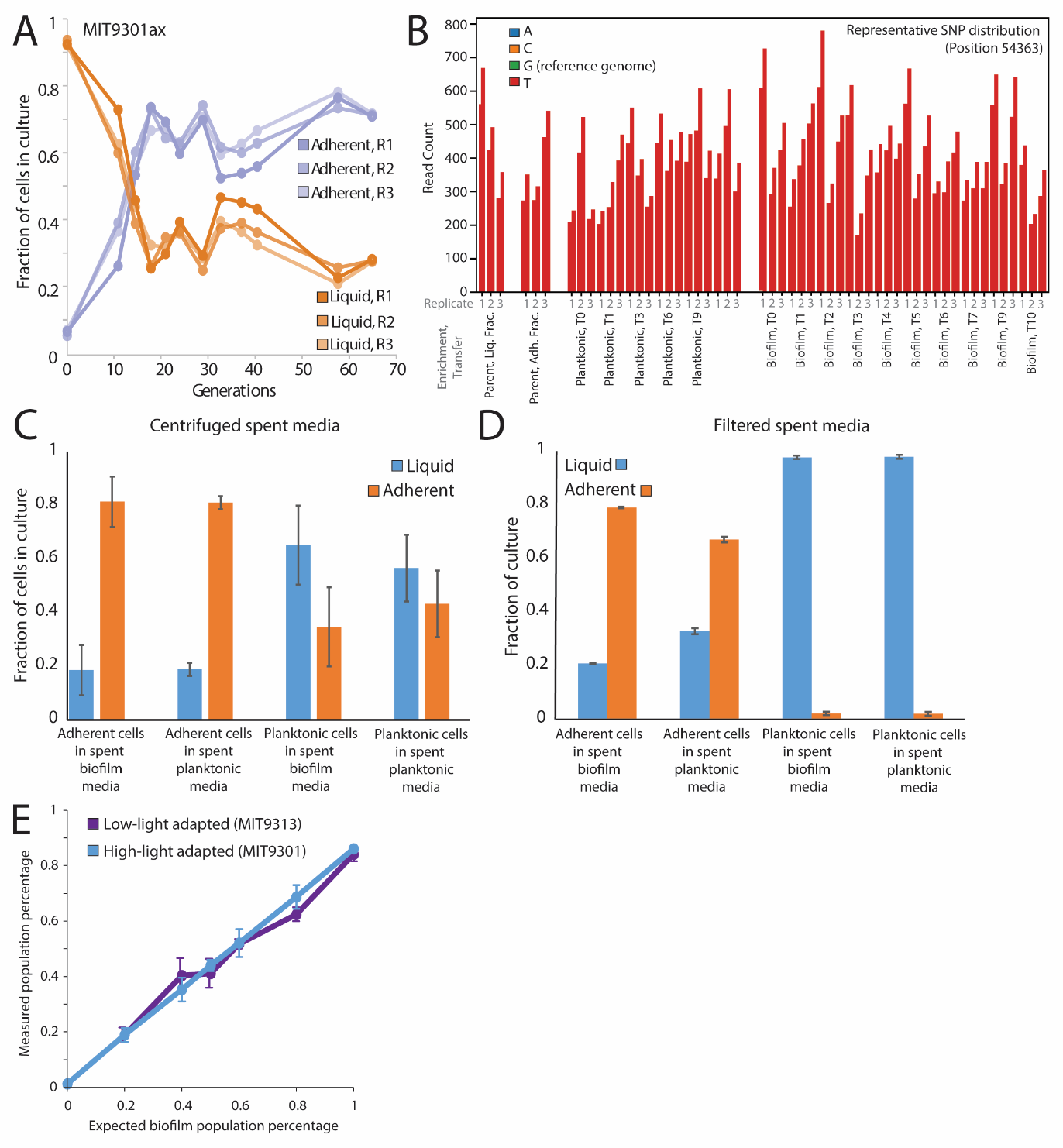

**Supplementary Figure 4. The PstS amino acid sequences from *Pseudomonas aeruginosa* and *Prochlorococcus* are in strong agreement.** Alignment of *Pseudomonas aeruginosa* (NP_254056.1) and *Prochlorococcus marinus* (WP_041484704.1) PstS amino acid sequence with an 87% query cover, 8x10^-7^ e-value, and 24.45% identity. Conservation indicates how closely the properties of the aligned amino acids match (top score 11 marked as *, score 10 marked as +). Quality is normalized from 0 to 1 (best) across the entire alignment, and then the consensus sequence and occupancy (identifying any alignment gaps) are calculated. (Visualized using Jalview)

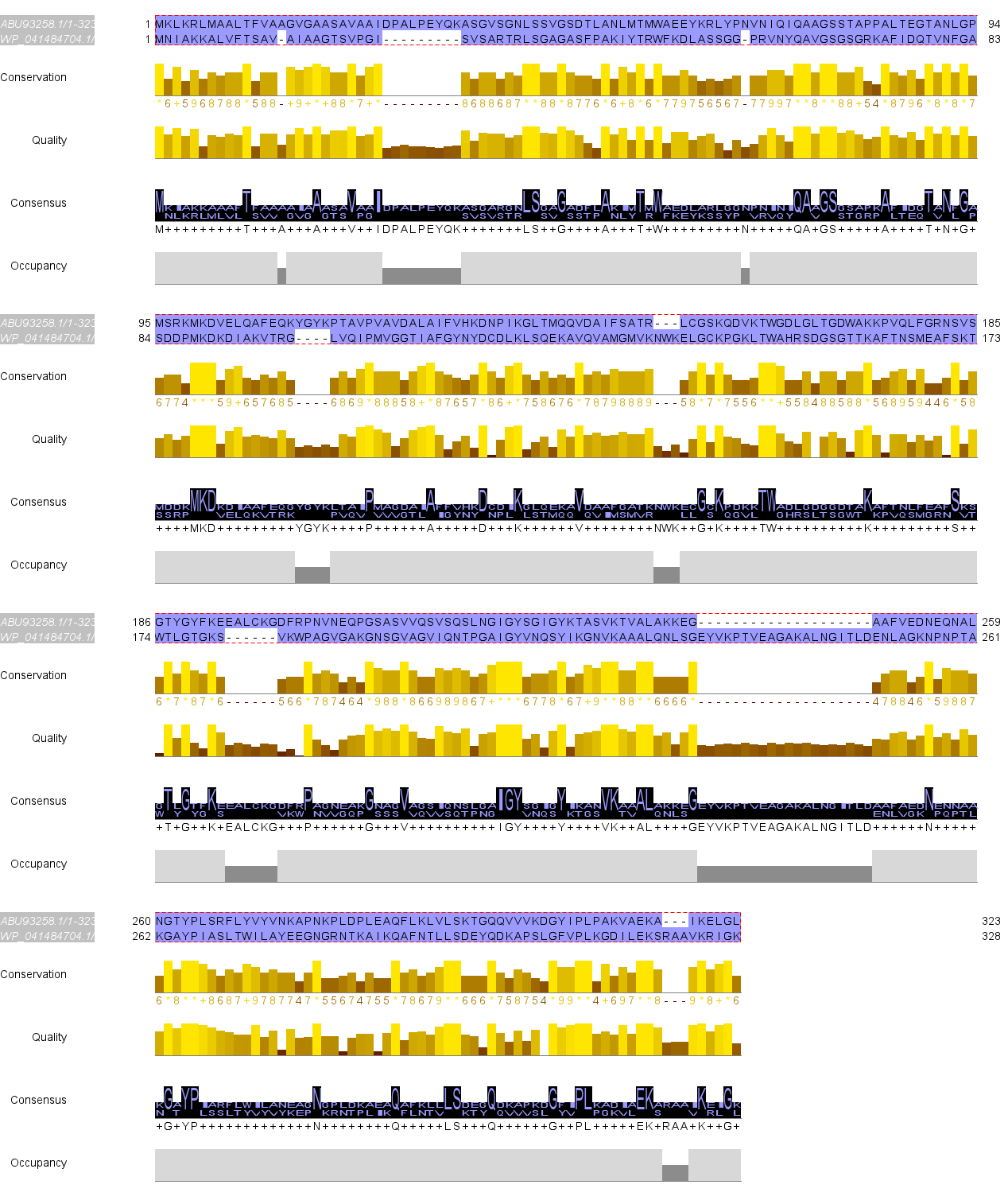

**Supplementary Figure 5. Chitin beads incubated with biofilm populations had approximately four times more attached cells than beads incubated with planktonic populations.** Chitin beads were incubated with MIT9301ax cells from either planktonic-enriched or biofilm-enriched cultures, and the cell concentrations in different culture fractions were quantified. First, the cell concentration in the media in each culture was measured (“initial liquid fraction”). At mid-exponential growth, planktonic cultures with chitin beads had 4.0x10^8^±7.0x10^6^ cells/mL in the liquid phase, while biofilm cultures were about 100-time less concentrated with 5.5x10^6^±3.1x10^5^ cells/mL (n=6). Next, the chitin beads were repeatedly washed with fresh media, and we measured the concentration of cells that washed off the beads in either culture (“washed off”). Slightly more cells were washed off chitin beads incubated with planktonic cultures (3.3x10^5^±1.5x10^4^ compared to 2.7x10^5^±3.6x10^3^, n=6). Finally, a subset of beads was vortexed for 3 minutes to dislodge all cells that remained attached to the chitin in either culture (“vortexed off”), and beads incubated with biofilm populations had a significantly higher number of attached cells compared to beads incubated with planktonic populations (9.9x10^4^±2.4x10^3^ and 2.5x10^4^±1.6x10^4^, respectively, n=3, p=0.05).

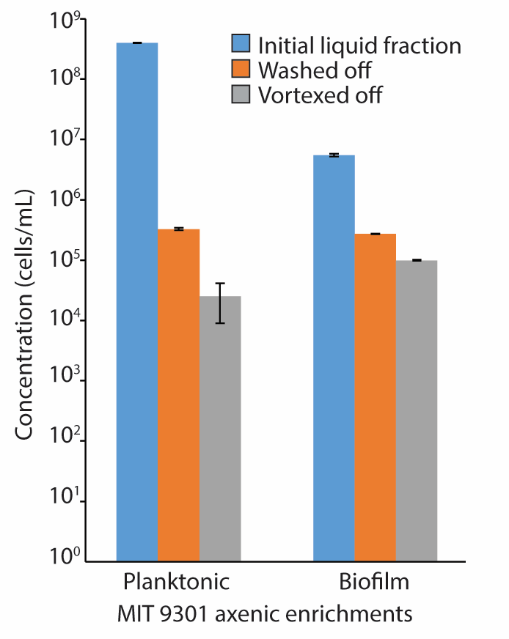
